# Vagus nerve stimulation modulates information representation of sustained activity in layer specific manner in the rat auditory cortex

**DOI:** 10.1101/2025.02.08.637217

**Authors:** Tomoyo Isoguchi Shiramatsu, Kenji Ibayashi, Kensuke Kawai, Hirokazu Takahashi

## Abstract

The brain of a living organism enables stable information processing in response to constantly changing external environments and internal states. As one of such cortical modulation, the present study focused on the effect of vagus nerve stimulation (VNS) therapy on information representation of the auditory cortex. By quantifying sound representation using machine learning, we investigated whether VNS alters cortical information representation in a layer-specific and frequency band-specific manner. A microelectrode array meticulously mapped the band-specific power and phase-locking value of sustained activities in every layer of the rat auditory cortex. Sparse logistic regression was used to decode the test frequency from these neural characteristics. The comparison of decoding accuracy before and after the application of VNS indicated that sound representation of the high-gamma band activity was impaired in the deeper layers, i.e., layers 5 and 6, while it was slightly improved in the superficial layers, i.e., layers 2, 3, and 4. Moreover, there was an improvement of sound representation in theta band activity in the deeper layers, demonstrating the layer-specific and frequency band-specific effect of VNS. Given that the cortical laminar structure and oscillatory activity in multiple frequency bands helps the auditory cortex to act as a hub for feed-forward and feed-back pathways in various information processing, the current findings support the possibility that VNS provide complex effects on brain function by altering the balance of cortical activity between layers and frequency bands.

## Introduction

The brain of a living organism is able to process sensory information in a stable manner by modulating the responsiveness of neurons and neural circuits according to the constantly changing external environment and internal state of the organism. For example, the neural responsiveness is dynamically adjusted according to the statistical properties of external stimuli, such as contrast and frequency of occurrence (Bonin et al., 2006; Lesica et al., 2007; Vinke et al., 2022). In addition, it has been reported that changes in the autonomic nervous system and neurotransmitters also regulate the transmission of sensory information processing (Noseda et al., 2017; Neves et al., 2018). Elucidating sustainable mechanisms for such modulation of neural circuits will undoubtedly contribute to the investigation of energy-efficient methods to bring flexibility and stability to artificial intelligence. As a such modulation of the neural circuit, which is deeply involved in changes in cognitive function, we have focused on cortical modulation induced by vagus nerve stimulation (VNS) therapy.

VNS has been acknowledged for its therapeutic effects, initially in refractory epilepsy (Zabara, 1992; Morris and Mueller, 1999; Ben-Menachem, 2002; Theodore and Fisher, 2004; Krahl and Clark, 2012) and, more recently, in depression (Rush et al., 2000; Wani et al., 2013), post-stroke rehabilitation (Engineer et al., 2019), and pain management (Schwedt and Vargas, 2015; Tassorelli et al., 2018). The clinical application of VNS in patients has revealed its modulatory effect on various brain functions, including memory, cognition, flexibility of thought, and creativity (Sackeim et al., 2001; Sjogren et al., 2002; Ghacibeh et al., 2006; Pena et al., 2014; Meisenhelter and Jobst, 2018). Based on the anatomical structure, it is widely acknowledged that VNS activates acetylcholine (ACh) (Detari et al., 1983; Clark et al., 1999; Hulsey et al., 2016; Collins et al., 2021; Mridha et al., 2021; Bowles et al., 2022), noradrenaline (NA) (Krahl et al., 1998; George et al., 2000; Groves et al., 2005; Hulsey et al., 2017), and serotonin (5-HT) (Rutecki, 1990; George et al., 2000; Ruffoli et al., 2011) systems through various neuronal nuclei. However, subsequent steps in altering brain functions, including VNS-induced modulation of information representation in the cerebral cortex, require further elucidation.

So far, we have made some progress in clarifying the laminar and oscillatory profiles of VNS-induced modulation on the auditory evoked responses in the rat auditory cortex (Takahashi et al., 2020; Kumagai et al., 2023). This has allowed us to accumulate electrophysiological evidence that VNS may provide pathway-dependent modulation of the brain, that is, different modulation of the feed-forward and feed-back pathways. The auditory cortex, with its laminar structure, along with other sensory cortices, has been suggested to function as a hub for these pathways, contributing to the various information processing (Linden and Schreiner, 2003; Markov et al., 2014). We have demonstrated that VNS enhances cortical activities relating to the feed-forward pathway, such as auditory evoked responses in superficial layers and gamma band oscillatory activities, while VNS diminishes those relating to the feed-back pathway, such as auditory evoked responses in deep layers and theta band oscillatory activities. It is hoped that this hypothesis of VNS-induced pathway-dependent neural modulation will be able to explain various outcomes of VNS in a unified manner, with demonstration that VNS also modulates neural representation in pathway-dependent manner.

Here, in the present study, our focus is on the laminar and oscillatory profiles of VNS-induced modulation on information representation in the sustained activity of the rat auditory cortex. It has been shown that a dense map of several characteristics in the sustained activity, such as band-specific power and phase locking value (PLV), can be decoded by machine learning and exhibits layer- and band-specific information representation(Shiramatsu et al., 2016b; Shiramatsu et al., 2019). he present study utilizes the same technique to investigate layer- and band-specific modulation on the information representation by comparing decoding accuracies before and after VNS was applied to the tested animals.

## Materials and Methods

This study adhered strictly to the “Guiding Principles for the Care and Use of Animals in the Field of Physiological Science,” published by the Japanese Physiological Society. The experimental protocol received approval from the Committee on the Ethics of Animal Experiments at the Research Center for Advanced Science and Technology at the University of Tokyo (Permit Number: RAC130107). All surgeries were performed under isoflurane anesthesia (3.5–4% v/v in the air for induction and 0.8–2.5% for maintenance), and meticulous efforts were taken to minimize suffering. Following the experiments, the animals were euthanized with an overdose of pentobarbital sodium (160 mg/kg, intraperitoneal administration).

### Implantation of the VNS system

Twenty-one male Wistar rats, aged 11–12 postnatal weeks and weighing 250– 350 g at the time of neural recording, were used in the experiments. The VNS system (VNS Therapy System Model 103, Cyberonics, TX, USA) was implanted in the rats under isoflurane anesthesia (Mylan Inc., PA, USA; 3.5–4% v/v in the air for induction and 0.8– 2.5% for maintenance) more than 4 days before the neural recording (Shiramatsu et al., 2016a; Takahashi et al., 2020; Kumagai et al., 2023). As shown in Fig. 1A, a skin incision was performed in the neck, and the spiral electrodes of the system were coiled around the exposed left vagus nerve. Each spiral electrode was 10 mm in diameter and 10 mm long, made of stainless steel, covered with polyurethane except where it contacted the vagus nerve. Following the subcutaneous implantation of the pulse generator on the back, the neck was sutured. Subsequently, the rats were allowed to recover from the implantation in their home cages with food and water. After the implantation and before the neural recordings described below, the impedance of the spiral electrode was confirmed to be sufficiently low (1000–2500 Ω, which is in the normal range of 600–5300 Ω.).

**Figure 1.**
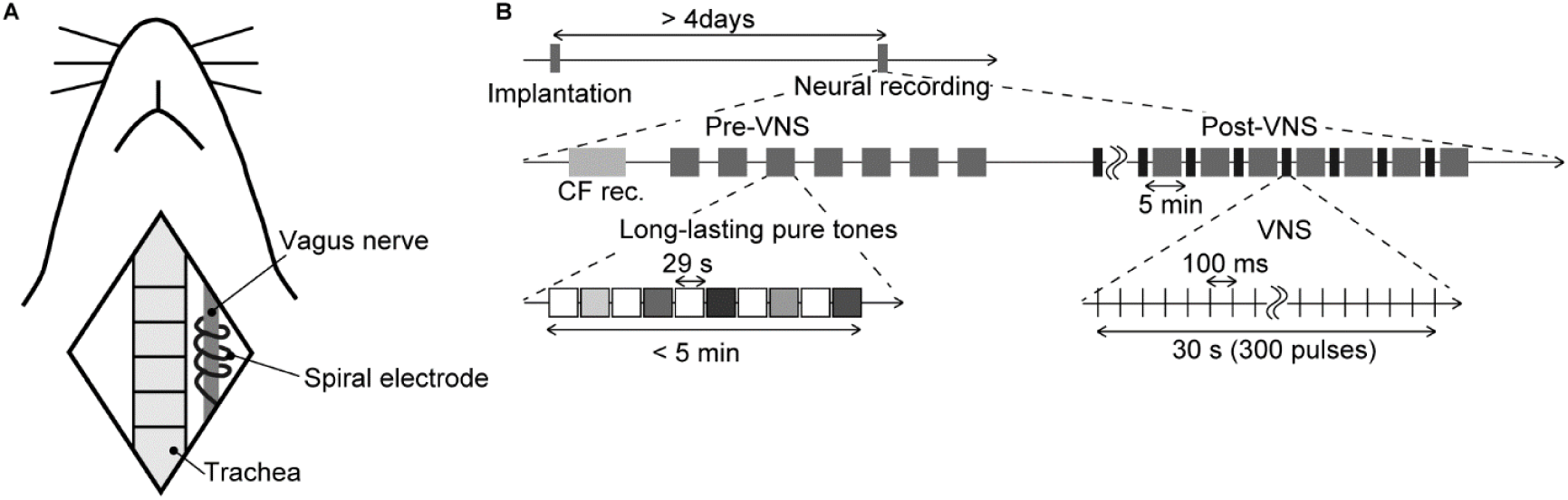
Schematic diagram of the experiment. (A) Animal preparation. The illustration shows anatomical landmarks of vagus nerve in a rat. The spiral electrodes of the system were coiled around the left vagus nerve. (B) Time course of the experiment. The tested animals were implanted with the stimulator more than 4 days before the neural recording. In the electrophysiological recording, we first performed the recording of the characteristic frequency (CF rec.) at each recording site, then two recording sessions, i.e., pre-VNS and post-VNS, were performed. In each session, we presented a sequence of long-lasting pure tones (29 s in duration, including a 5-ms rise/fall, 60 dB SPL). Test frequencies were 8.0, 10, 13, 16, and 32 kHz, and were randomized across the seven sequences. Each sequence did not exceed 5 minutes. In the post-VNS session, VNS was applied with a current of 500 μA, a pulse width of 130 μs, and a stimulation frequency of 10 Hz. The system stimulated for 30 s (300 pulses), alternating with a 5-min rest, during which the sequence of long-lasting pure tones was presented again.

### Neural recordings

A second surgery was performed to conduct electrophysiological recordings from each layer of the auditory cortex. The implantation procedure of the microelectrodes followed the same protocol as detailed in the previously published papers (Shiramatsu et al., 2016b; Shiramatsu et al., 2019; Kumagai et al., 2023). The rats were once again anesthetized with isoflurane and secured in place using a custom-made head-holding device. Atropine sulfate (0.1 mg/kg; Nipro ES Pharma Co., Ltd., Osaka, Japan) was administered at the surgery’s commencement and conclusion to diminish bronchial secretions’ viscosity. A skin incision was made at the commencement of the surgery under local anesthesia using xylocaine (1%, 0.1–0.5 mL; Aspen Japan, Tokyo, Japan). The right temporal muscle, cranium, and dura covering the auditory cortex were excised surgically. The exposed cortical surface was perfused with saline to prevent desiccation, and the cisternal cerebrospinal fluid was drained to reduce cerebral edema. A needle electrode was subcutaneously inserted into the right forepaw and served as a ground. Near the bregma landmark, a small craniotomy was performed to embed a 0.5-mm-thick integrated circuit socket as a reference electrode, establishing electrical contact with the dura mater. The right eardrum ipsilateral to the exposed cortex was intentionally ruptured and sealed with wax to ensure unilateral sound input from the ear contralateral to the exposed cortex.

A heating blanket was utilized to sustain the body temperature at approximately 37 °C. Throughout the experiment, respiration rate, heart rate, and hind paw withdrawal reflexes were monitored to ensure an adequate and stable level of anesthesia.

Consistent with our previous study, a microelectrode array (ICS-96, Blackrock Microsystems, Salt Lake City, UT, USA) with a 10 × 10 grid of recording sites within a 4 × 4 mm area simultaneously recorded local field potentials (LFPs) from layer 2/3 (L2/3, *n* = 7), layer 4 (L4, *n* = 7), or layer 5/6 (L5/6, *n* = 8) of the auditory cortex at depths of 350, 700, or 1,000-μm, respectively. The four recording sites at the corners of the grid were offline, and 96 recording sites were utilized for the recordings. LFPs were acquired with an amplification gain of 1,000, a digital filter bandpass of 0.3–500 Hz, and a sampling frequency of 1 kHz (Cerebus Data Acquisition System, Cyberkinetics Inc., Salt Lake City, UT, USA). Acoustic stimuli were generated using a function generator (WF1947, NF Corp., Kanagawa, Japan), and a speaker (Technics EAS-10TH800, Matsushita Electric Industrial Co. Ltd., Osaka, Japan) was positioned 10 cm from the left ear (contralateral to the exposed cortex). Test stimuli were calibrated to 60 dB SPL (concerning 20 μPa) at the pinna using a 1/4-inch microphone (Type 4939, Brüel & Kjaer, Nærum, Denmark) and a spectrum analyzer (CF-5210, Ono Sokki Co., Ltd., Kanagawa, Japan).

Once the injury spike was confirmed to have subsided, two sessions of neural recordings–– pre-VNS and post-VNS recordings, were carried out (Fig. 1B). In each session, we recorded LFPs as sustained activities in response to long-lasting pure tones (29 s in duration, including a 5-ms rise/fall, 60 dB SPL), covering frequencies from 8.0– 32 kHz (8.0, 10, 13, 16, and 32 kHz). Each pure tone was repeated seven times in a pseudorandom order, interleaved with 29-s silent blocks. Following the initial (pre-VNS) recording, we commenced the application of VNS. The electrical pulses for VNS were biphasic and charge-balanced to prevent damage to nerve fibers. The first and second phases featured short-term high and long-term low amplitudes designed to activate afferent fibers. In the first phase, the current was set at 500 μA with a pulse width of 130 μs, and the second phase had a lower current and a longer width than the first phase, to balance the total charge but not to stimulate efferent fibers in the vagus nerve. The VNS system was stimulated at a stimulation frequency of 10 Hz for 30 s, a total of 300 pulses, alternating with a 5 min of rest. The second (post-VNS) recording was conducted after more than seven instances of VNS had been applied, which was more than 30 min from the initial application of VNS. During the second recording, long-lasting pure tones were presented during a 5-min resting interval to prevent recording neural activities directly induced by VNS. Each session of neural recordings took approximately 40 min.

### Decoding of test frequency

Data analysis was performed using MATLAB (MathWorks, Natick, MA, USA). Similar to our previous studies (Shiramatsu et al., 2016b; Shiramatsu et al., 2019), machine learning, such as sparse logistic regression (SLR), decoded the test frequency from the band-specific power or PLV in one of five frequency bands (theta, 4–8 Hz; alpha, 8–14 Hz; beta, 14–30 Hz; low-gamma, 30–40 Hz; and high-gamma, 60–80 Hz). Subsequently, the decoding accuracy using each neural characteristic was compared between the pre- and post-VNS conditions. This study follows previous studies and set a relatively narrow frequency bandwidth for each frequency band in order to properly calculate instantaneous angles with the Hilbert transform in the following step.

Band-specific power and PLV were initially extracted from sustained LFPs in response to long-lasting pure tones. Bursting LFPs, which manifest under isoflurane anesthesia regardless of sound presentation and consequently hinder the decoding of sound information, were eliminated as previously described (Shiramatsu et al., 2016b; Shiramatsu et al., 2019). Bursting LFPs were classified when the standard deviation for each 100-ms LFP exceeded the threshold at more than 24 recording sites (Fig. 2A and B). Subsequently, band-specific power and PLV were calculated from 1,000-ms continuous non-burst waves (Fig. 2A). The band-specific power at each recording site was calculated as the root mean square of the bandpass-filtered LFPs. The PLVs between all 4,560 pairs of the 96 recording sites were calculated using the following equation:

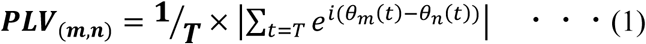

**Figure 2.**
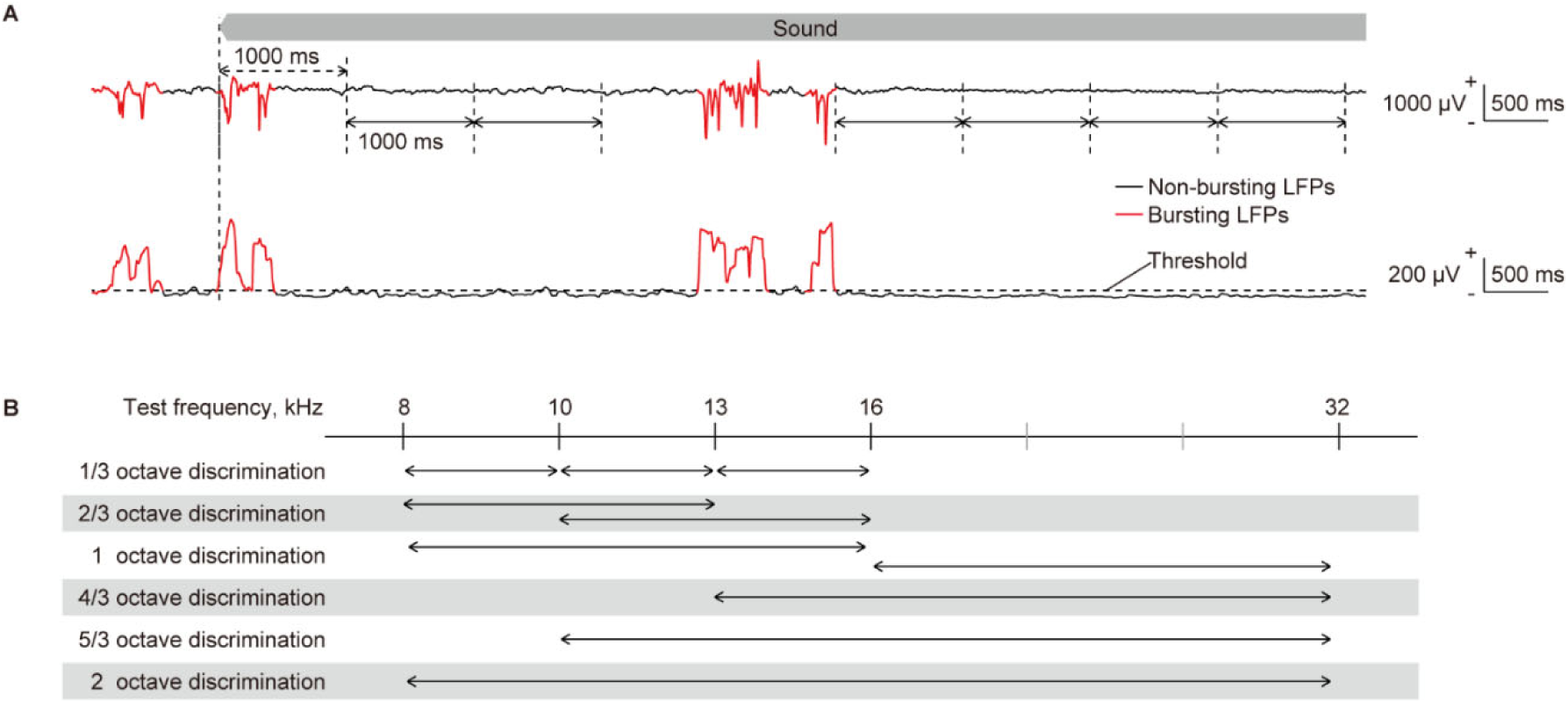
Quantification of neural characteristics and decoding of test frequency. (A) (top) Representative raw traces of sustained local field potentials (LFPs) recorded from layer 2/3 in response to a pure tone of 16 kHz. LFPs during the initial second of sound presentation were considered as transient onset activity, as indicated by double-headed arrows with dashed lines. They were excluded from the quantification of neural characteristics. Following the onset response, the LFPs were separated by one consecutive second, as indicated by double-headed arrows with solid lines, from each of which band-specific power and PLV were calculated. The red and black traces of LFPs represent bursting and non-bursting LFPs, respectively, that were classified (please see the method section and the following figure). (bottom) the standard deviation (SD) of 100-ms LFPs, including burst activities, is shown. Time intervals where the SD exceeds a threshold in more than 24 recording sites and persists for durations of 150 ms or longer were classified as bursting LFPs (red line), while others below these criteria were classified as non-bursting LFPs (black line). (B) Two-choice frequency decoding was conducted using frequency pairs with various frequency ratios. Given that we presented pure tones of five frequencies, we paired two. We applied six different decoding with distinct frequency distances, representing varying difficulty levels of distinction.

Here, *m* and *n* denote the recording site numbers, θ represents the instantaneous angle at each time point obtained by the Hilbert transform of the filtered LFP, *T* indicates the time included in the 1,000-ms waves, and *i* is an imaginary unit. The PLV is a real value ranging from 0–1, signifying that the band-specific activities at the two recording sites were perfectly desynchronized and synchronized. During each recording session, i.e., pre- and post-VNS sessions, we acquired 70 patterns of band-specific power and PLV for each test frequency. These patterns served as the datasets for SLR.

To scrutinize sound representation in the auditory cortex, we performed two types of decoding based on the patterns of each neural characteristic. The first involved decoding of the five test frequencies, while the second focused on the two-choice discrimination of two among the five test frequencies (Fig. 2B). All discriminations were conducted independently for each tested animal, each frequency band for the LFP, and each recording session (e.g., pre- and post-VNS), according to the following procedure using SLR toolbox ver. 1.2.1 alpha as a toolbox for MATLAB (Miyawaki et al., 2008; Yamashita et al., 2008).

Details of the decoding process have been previously described (Shiramatsu et al., 2019). Briefly, the input data for the SLR were labeled using the test frequencies. Initially, the 70-input data for each label were separated into 60 for supervised learning and 10 for accuracy testing. Supervised learning was then applied to these combinations of data and labels, utilizing 300 or 120 data in the five- or two-choice discrimination, respectively. After the supervised learning, SLR discriminated the novel test data (10 for each label), and we calculated the percentage of successfully discriminated data as the accuracy rate for decoding. Seven-fold cross-validation was employed, and ultimately, the mean accuracy rates for all cross-validations and test frequencies were calculated for each animal.

As a statistical test to examine whether VNS affects sound representation for each neural characteristic, the accuracy rate in the pre-VNS session was subtracted from that in the post-VNS session. Each change in the accuracy rate was then compared with zero using a two-sided Wilcoxon signed-rank test, with 0.05 adapted as the significance level for the p-value.

## Results

Figure 3 depicts the representative patterns of the band-specific power and PLV in the high-gamma band in each layer. As discussed in previous studies, sustained activities demonstrate ambiguous patterns compared to onset activities, such as auditory evoked responses (deCharms and Merzenich, 1996; Eggermont, 1997; Takahashi et al., 2005; Polley et al., 2007; Shiramatsu et al., 2016b). In response to low test frequency tones, two foci band-specific powers were observed in the posterior-dorsal and anterior-ventral parts of the auditory cortex. Meanwhile, there was a single focus at the anterior-medial part of the auditory cortex for higher test frequency tones. Consistent with the findings in the previous study, SLR could decode the test frequencies from these patterns, with a high accuracy rate observed in high-gamma bands (Shiramatsu et al., 2016b; Shiramatsu et al., 2019).

**Figure 3.**
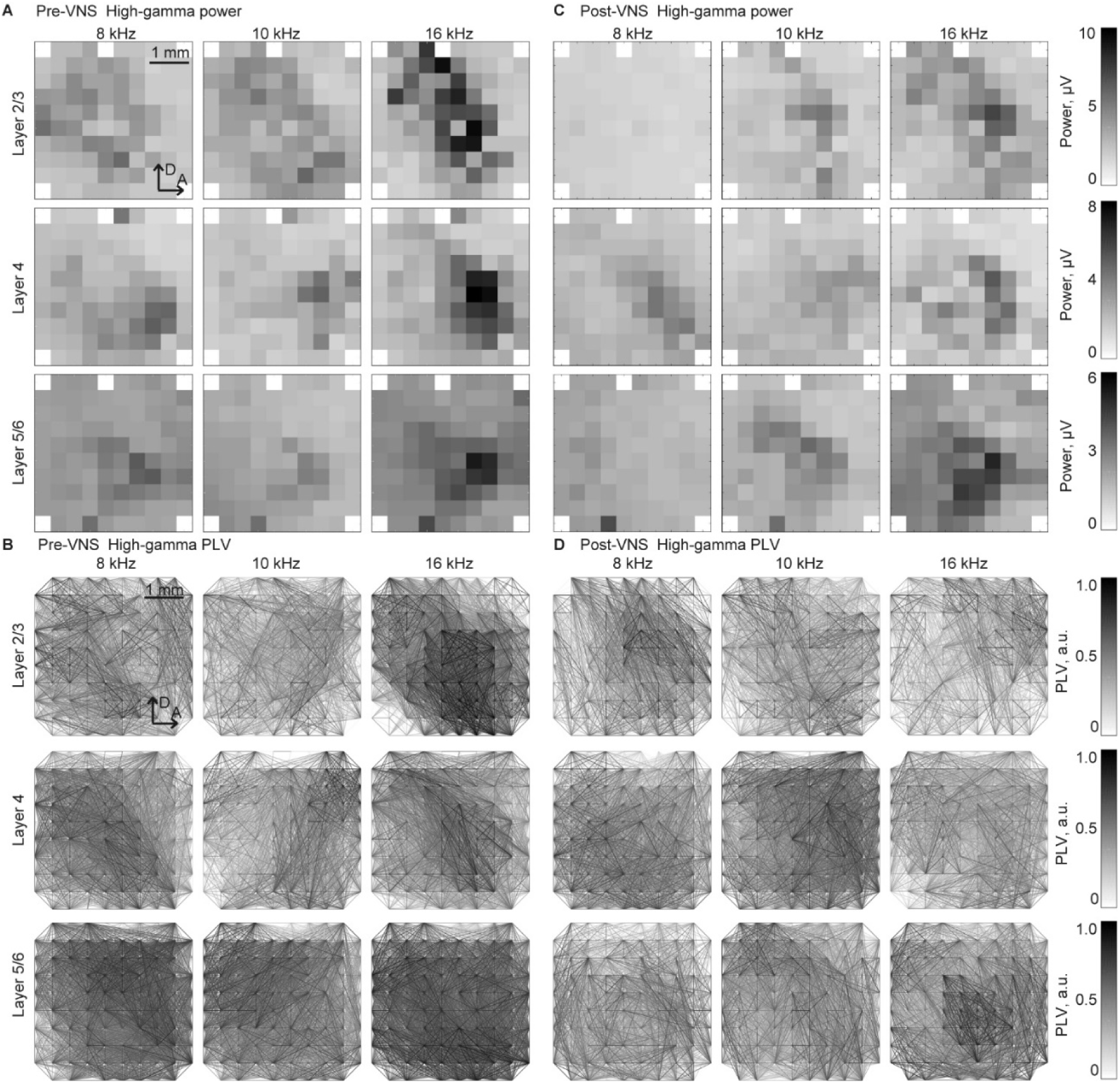
Spatial patterns of the band-specific power and phase locking value (PLV) for the frequency decoding Representative spatial maps of (A, B) band-specific power and (C, D) phase-locking value (PLV) in the high-gamma band in response to selected test frequencies, 8, 10, and 16 kHz. The spatial maps for the recording from layers 2/3 (top), 4 (middle), and 5/6 (bottom) are displayed in three representative animals. The patterns exhibited similarities between the recordings (A, C) before and (B, D) after the application of VNS. The tonotopic shifts of the response foci are well identified, particularly in the spatial maps of the band-specific power.

The point of interest in this study was the VNS-induced changes in decoding accuracy. Consequently, the accuracy rate in the pre-VNS session was subtracted from that in the post-VNS session (Fig. 4). In the five-choice test frequency decoding, several significant changes were observed at L5/6. The decoding accuracy in this layer significantly improved in band-specific power in the theta band while impaired in the PLV in the high-gamma band (Fig. 4A and B, Wilcoxon two-sided signed-rank test versus 0%, *p* = 0.047 and 0.031). No significant changes were observed in the other layers.

**Figure 4.**
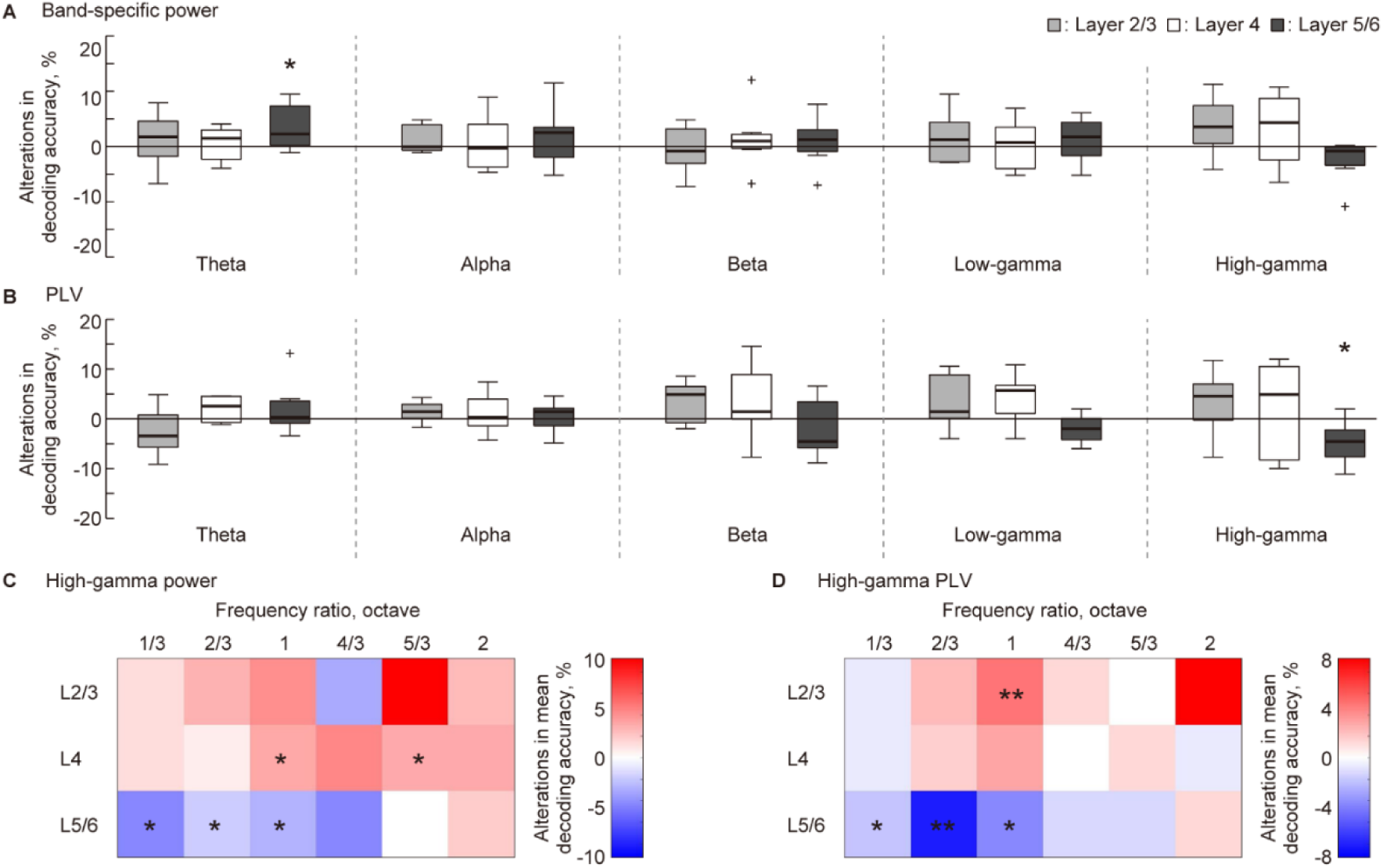
Layer-specific VNS-induced changes in the decoding accuracy of sparse logistic regression. (A, B) In the five-choice discrimination of the test frequency, decoding accuracy for pre-VNS activities was subtracted from that of post-VNS activities. The alterations in decoding accuracy for each frequency band of sustained activity and each layer was accessed. Statistical tests for the change in decoding accuracy in (A) band-specific power and (B) phase locking value (PLV) were conducted. (C, D) We obtained the difference between median decoding accuracy in pre- and post-VNS recordings in the two-choice discrimination across six frequency ratios. The red and blue density scales represent the improvement and inhibition of the discrimination, respectively. Asterisks indicate that the median of the changes in decoding accuracy is significantly higher or lower than zero: * *p* < 0.05, ** *p* < 0.01 (Wilcoxon rank sum test).

A previous study indicated that sustained activity in the auditory cortex most effectively represents the test frequency in the high-gamma band (Shiramatsu et al., 2016b; Shiramatsu et al., 2019). Based on this, we delved into the decoding frequency resolution of decoding in the high-gamma band through two-choice discrimination of test frequency at six distinct frequency ratios (Fig. 2C). Figure 4 (C–D) illustrates the VNS-induced median change in discrimination accuracy. The red and blue density scales signify improvement and inhibition of discrimination, respectively. In L5/6, discrimination between close frequencies, especially those ≤ one octave, tended to be inhibited (Wilcoxon two-sided signed-rank test versus 0%, Fig. 4C, p = 0.035, 0.025, and 0.012; Fig. 4D, p = 0.027, 0.0040, and 0.033, respectively). Conversely, in L2/3 and L4, the median discrimination accuracy often increased, with some instances being significant (Fig. 4C, p = 0.025 and 0.039 for L4; Fig. 4D, p = 0.015 for L2/3).

## Discussion

The present hypothesis was that VNS alters cortical information representation in the auditory cortex in a layer-specific and frequency band-specific manner. The SLR performed several decoding and discrimination tasks of the test frequency from the band-specific power and PLV calculated from the sustained activities. Comparing the differences in decoding accuracy revealed that VNS mainly impaired sound representation in L5/6, while it slightly improved sound representation in L2/3 and L4. These results mark the first demonstration of VNS-induced functional changes in the sensory cortex and corroborate the earlier suggestion of the layer-specific cortical effect of VNS.

### VNS-induced acute modulation of sound representation in the auditory cortex

By focusing on sustained activity, this study demonstrates that VNS has an acute effect on the cortical representation of sound frequency, a principal function of the auditory cortex. In each sensory modality, the sensory cortex represents the physical characteristics of the sensory input. In the auditory cortex, the tonotopic map generates distinct spatial patterns of neural activity corresponding to different sound frequencies (Takahashi et al., 2005; Polley et al., 2007). Dense mapping with microelectrode arrays provides tonotopic spatial patterns, and machine learning can easily decode the test frequencies (Funamizu et al., 2011; Shiramatsu et al., 2016b). This tonotopic organization is primarily established by the hard-wired feed-forward pathway from the cochlea, where the asymmetric structure produces the initial tonotopically separated patterns (Merzenich et al., 1975; Dallos, 1996; Malmierca, 2003; Mann and Kelley, 2011; Tobin et al., 2019). Conversely, cortical tonotopy changes plastically through associative learning and exposure to the acoustic environment during critical periods (Bakin and Weinberger, 1990; Recanzone et al., 1993; de Villers-Sidani et al., 2007; Mann and Kelley, 2011). Therefore, the hard-wired yet plastic tonotopic organization and its representation of sound frequency are considered one of the most fundamental functions of the auditory cortex.

The tonotopic structure of the auditory system is defined based on the transient onset activity such as the auditory evoked response that promptly follows the onset of sounds(Doron et al., 2002; Rutkowski et al., 2003; Takahashi et al., 2005; Polley et al., 2007). Owing to the high reproducibility and decodability mentioned earlier, transient activity may not be suitable for illustrating VNS effects, which are not expected to be powerful enough to change the tonotopic spatial pattern. For instance, associative learning-induced changes in tonotopy often require hours to manifest (Ide et al., 2012). Moreover, altering the characteristic frequency of each neuron organizing the cortical tonotopy requires paired stimulation of VNS and specific sounds (Engineer et al., 2013; Borland et al., 2016). As in this study, previous attempts found it difficult to change the sound representation of transient activities by one-hour VNS without pairing. A similar examination using the same approach as this study achieved almost 100% decoding accuracy from the transient activity in pre- and post-VNS conditions without any change (data not shown). However, sustained activity is less reproducible and, therefore, less decodable than transient activity because sound-induced activity quickly attenuates within several hundred milliseconds (Smith and Zwislocki, 1975; Smith, 1979; Westerman and Smith, 1984). Neuromodulations can easily influence this ambiguous sound representation. Therefore, machine learning in the present study successfully quantified the effect of VNS.

The information representation of sustained activity showed robust VNS-induced change in the high-gamma band. Consistent with findings from a previous study, the information representation of sustained activity is contingent on the cortical layer and frequency band of neural activity (Shiramatsu et al., 2019). Notably, the decoding accuracy of the band-specific power and PLV was better in the high-gamma bands in L4 and L5/6 (approximately 50% for five frequency discriminations, with a chance level of 20%). In contrast, it was only slightly (yet significantly) above the chance level in the other bands in these layers. Additionally, the decoding accuracy in L2/3 was low for all the bands, indicating that the information representation in the supragranular layer is inherently weak (Winkowski and Kanold, 2013). Considering that inhibitory cells in the deeper layer of the auditory cortex, responsible for generating high-gamma oscillations and mediating lateral inhibition, are primary candidates for the sound representation of sustained activity (Jefferys et al., 1996; Hasenstaub et al., 2005; Bartos et al., 2007), this study values changes in decoding accuracy in the high-gamma band. Significant improvements in decoding accuracy were noted for band-specific power in the theta band in L5/6 (Fig. 3A); however, it is worth noting that the decoding accuracy was relatively low in the pre-VNS condition, and the improvement was still modest. Additionally, the subsequent two-choice discrimination of test frequencies revealed no consistent or robust changes in decoding accuracy in this band and layer (data not shown). Consequently, this study values the improvement in the high-gamma band in L5/6, indicating that the effect of VNS on cortical sound representation is mainly mediated by changes in the high-gamma oscillation in the deeper layers, such as lateral inhibition.

### Possible mechanism

The current findings regarding the effects of VNS on auditory cortex function consistently support the previous hypothesis of neuromodulatory effects of VNS on the sensory cortex through several neurotransmitters. First, the VNS-induced changes in the information representation of sustained activity reported in this study were akin to the changes in the magnitude of AEP reported in a previous study, showing a similar layer dependence (Takahashi et al., 2020). The previous study reported that the VNS-induced enhancement of AEP was more predominant in superficial layers than in deeper layers. AEP most strongly reflects feed-forward neurotransmission from the auditory periphery (Linden and Schreiner, 2003; Malmierca, 2003; Lee and Winer, 2008); a similar facilitation of feed-forward neurotransmission also enhances sustained activities in the superficial layers, thereby making the sound representation more robust in these layers. This aligns with the several improvements in decoding accuracy observed in the high-gamma band in L2/3 and L4. Additionally, the tendency of the VNS effects to be opposite between the superficial and deep layers was also consistent with the changes in decoding accuracy, improvement, and impairment, respectively.

Second, based on previous surface recordings, it was predicted that changes in gamma-band oscillations would be predominant and opposite to those in the low-frequency band(Kumagai et al., 2023). AEP obtained by ECoG recording demonstrated that a 2-h application of VNS enhanced and diminished the sound-induced oscillatory power in the gamma and theta bands, respectively. It remains unclear which cortical layer is primarily reflected by the oscillatory activity at the cortical surface, and there were still two commonalities with the present results: a dominant change in the high-frequency oscillation and an opposite effect of VNS on higher and lower frequency oscillations. Our results substantiate a layer-specific mechanism of the effect of VNS on the auditory cortex, consistent with previous studies.

Currently, the best explanation for the layer-specific effects of VNS is the layer-specific distribution of neurotransmitter receptors in the sensory cortex. VNS activates the ACh, NA, and 5-HT systems through the basal forebrain, locus coeruleus, and raphe nucleus. Receptors for these neurotransmitters were distributed heterogeneously from superficial to deep cortical layers, consistent with the layer-specific effects of VNS observed in this study. Considering that a few hours are insufficient for a full-scale release of 5-HT, ACh, and NA seem to mediate the present effect of VNS. Previous pharmacological investigations with antagonists have demonstrated that VNS acts on the auditory cortex in gamma- and beta-oscillations through nicotinic receptors and in the theta-oscillation through NA, respectively. This aligns with the demonstrated effect of VNS on sound representation in the auditory cortex, which was strongest in the high-gamma band and significant in the theta band.

The altered information representation in the high-gamma and theta bands is consistent with the reported outcomes of VNS. The high-gamma band brought by cortical local interneuron mainly mediates intracortical information processing such as sparse coding in sensory cortex, as well as feature binding in visual perception. Such binding function of the gamma activity within sensory-cognitive processes has been suggested to include associative memory, as well as creativity. On the other hand, the theta band mainly mediates global intercortical connectivity to coordinate multiple brain regions. It is also suggested that such theta-band mediated global communication contributes to several types of learning and to epileptic network in its patients. Taken together, the present results made a first putative link between the cortical modulation and the functional outcomes of VNS, through the frequency band-specific effect of VNS.

The present study raises the possibility to explain the cortical modulation of VNS through alterations in the balance between cortical feed-forward and feed-back pathways. In the auditory cortex, which acts as the hub of the feed-forward and feed-back pathways, ACh controls sensory gating along the feed-forward pathway, and NA provides top-down feed-back control. Modulations of these neurotransmitters by VNS enhance and suppress the feed-forward and feed-back pathways, altering the balance between cortical feed-forward and feed-back. In the layered structure of the auditory cortex, L2/3, and L4 are on the feed-forward pathway, and L5/6 is responsible for the feed-back pathway. Therefore, the present results, indicating that VNS clarified sound representation in the former layers and made it ambiguous in the latter, strongly support the possibility that VNS influences brain function by altering the balance of the feed-forward and feed-back pathways.

### Methodological considerations and future directions

In the present study, the reference electrode was placed in the left parietal cortex, which was contralateral to the recorded right auditory cortex. Different placement of the recording electrode affects the recorded neural signals and sometimes reverses the polarity of a specific evoked responses. Moreover, large distance between the recording sites and the reference electrode is more likely to introduce global artifacts into the recorded signal. In the present recording, we did not consider such a global effect on the decoding accuracy, however, we believe that such an effect of the reference on the VNS-induced change was not too severe for the following two reasons. First, the present characteristics for the decoding were obtained from the sustained or steady-state activity, which were not directly affected by the reversal in instantaneous phase polarity. Even if the phase of a particular frequency band is inverted at a given time, it is canceled out by the root-mean-square process to obtain the band-specific power, and by the subtracting the instantaneous phase between the recording sites to obtain PLV. The effect of the reference positioning should be considered in further studies.

The study results should be considered in light of the effects of isoflurane anesthesia, as well as the time lapse between the pre- and post-VNS sessions. Anesthesia generally suppresses and inhibits excitatory and inhibitory synapses. Specifically, reports have indicated that isoflurane anesthesia suppresses excitatory NMDA receptors, enhances inhibitory GABAA receptors, and inhibits the feed-back pathway. Given that the sound representation in the pre-VNS and the impact of VNS itself may be affected by anesthesia, certain changes may have been exaggerated or overlooked in this study. Moreover, isoflurane anesthesia introduces burst inhibition, which divides neural activity into two states: intermittent bursting LFP, characterized by high-amplitude synchronization across the auditory cortex, and low-amplitude non-bursting LFP. It has already been demonstrated that band-specific power and PLV in the non-burst LFP encode sound frequency in each layer. In addition, the time lapse between the pre- and post-VNS sessions should be considered, although its effect on the present result may be small for the following two reasons. First, a previous ECoG recording, which made the same recording from the control animals not applied VNS, demonstrated that the oscillatory power in the auditory evoked response is stable for 2 hours. Second, even if there were a global change in the sustained activity over the time, it would be cancelled out by the machine learning discrimination applied separately to the pre- and post-VNS sessions. Taken together, the present study provided a first demonstration of the layer and frequency band-specific effect of VNS, however, further attempts should be made under alternative anesthetics or in the awake state and with sham stimulation or without VNS to fully reveal the effect of VNS.

To further explain the effects of VNS, similar studies should be conducted in the future on animal models of epilepsy and chronic VNS. In a previous study, VNS enhanced PLV in naive rats but weakened PLV in epileptic model rats already exhibiting strong synchrony. In this study, the compromised sound representation in L5/6 implies a decline in the feed-back pathway, regarded as a significant pathway in epilepsy propagation (although it is still controversial), especially in temporal lobe epilepsy. Subsequent demonstrations in animal models of epilepsy are expected to confirm the indication that VNS modulation on L5/6 contribute to seizure suppression. Furthermore, this study demonstrated the modulation of information representation by a few-hour VNS application, characterized as an acute effect. Considering prior reports on the clinical benefits of chronic VNS, including seizure suppression and learning enhancement, it would be valuable to investigate the impact of chronic VNS, i.e., from several days to months—on sound representation in the sensory cortex in the future.

This study is the first report indicating that VNS affects the function of the sensory cortex by altering the representation of sound information within a few hours in a layer-specific manner. This study further connects actual behavioral changes, such as memory learning and information processing in the sensory cortex—the feed-forward and feed-back pathways hub. Simultaneously, it also calls attention to additional studies of how this change in information representation affects memory learning and how it is mediated by microscopic neural mechanisms, i.e., the receptive fields in the auditory cortex, following the previous series of studies examining plastic changes induced by paired stimulation of VNS and sounds.

## Author Contributions

TIS: Conceptualization, Formal analysis, Funding acquisition, Methodology, Visualization, Writing – original draft; KI: Investigation, Methodology, Writing – review and editing; KK: Conceptualization, Funding acquisition, Resources, Writing – review and editing; HT: Conceptualization, Funding acquisition, Writing – review and editing.

